# Growth kinetics and power laws indicate distinct mechanisms of cell-cell interactions in the aggregation process

**DOI:** 10.1101/2021.12.22.473802

**Authors:** Debangana Mukhopadhyay, Rumi De

## Abstract

Cellular aggregation is a complex process orchestrated by various kinds of interactions depending on its environments. Different interactions give rise to different pathways of cellular rearrangement and the development of specialized tissues. To distinguish the underlying mechanisms, in this theoretical work, we investigate the spontaneous emergence of tissue patterns from an ensemble of single cells on a substrate following three leading pathways of cell-cell interactions, namely, direct cell adhesion contacts, matrix mediated mechanical interaction, and chemical signalling. Our analysis shows that the growth kinetics of the aggregation process is distinctly different for each pathway and bears the signature of the specific cell-cell interactions. Interestingly, we find that the average domain size and the mass of the clusters exhibit a power law growth in time under certain interaction mechanisms hitherto unexplored. Further, as observed in experiments, the cluster size distribution can be characterized by stretched exponential functions showing distinct cellular organization processes.

## INTRODUCTION

One of the most fundamental aspects of developmental biology is the ability of cells to aggregate and form tissues. Understanding the underlying mechanisms of cell-cell interactions in the aggregation process and the consequent tissue formation is immensely important to design therapeutic approaches for a number of diseases, cancer metastasis, wound healing, tissue regeneration, and in diverse areas in developmental biology [1–4]. Several studies have been carried out to unravel the self-assembly and organization process in both living as well as non-living systems, such as flocking of birds [5], the formation of diverse swarms [6–8], patterns of bacterial colony [9], aggregation of proteins [10], assembly of active filaments [11], growth of intricate snowflakes [12] among many others. In such collective systems, the interactions among the constituent entities have been found to play a crucial role in determining the emergence of self-organized patterns. Biological cells, however, live in very complex physiological environments. Nature of the cellular interactions strongly depend on its surroundings and also varies with the diverse processes that it performs. Cellular organizations take place through various pathways via direct cell adhesion contacts, mechanical interactions mediated through the extracellular matrix, or chemical signalling [13–15]. Each of these mechanisms provides a distinctly different route for cell-cell communications leading to cell-assembly and unique tissue structures.

Multiple experimental and theoretical approaches have been developed to probe the signals and characterize the underlying mechanisms of the cellular aggregation process in living tissues [2, 3, 16–19]. Salm and Pismen have studied the long-range coordination of multiple cells by chemical and mechanical signal in epithelial spreading [18]. They have shown that the feedback between cell deformation and intracellular signal transduction plays a key role in maintaining a directional collective cell migration. Studies have further shown, some cell types spontaneously reorganize to increase the cell cohesivity through the adhesion junction proteins to give rise to various morphological structures. These observations have been elucidated by the differential adhesion hypothesis (DAH), which approaches the problem from a simple physical perspective that motile cohesive cells tend to assemble into aggregates to achieve minimal interfacial energy and maximal binding strength [20, 21]. Song *et al.* have experimentally shown that human embryonic stem cells-derived pancreatic progenitors are purified from co-cultured feeder cells using this spontaneous self-organization. During the cell growth process, low cohesive cells surround the high cohesive cells on the periphery [22]. DAH could explain a wide range of cellular behaviours such as cell sorting, aggregation, spreading of tissues, and tumour growth [20, 21, 23]. Moreover, it is found that cell assembly is also strongly sensitive to the substrate stiffness [24]. Using substrates of identical chemical composition but varying the rigidity, one observes the formation of different tissue patterns in the cell cultures. On a stiff substrate, cells migrate away from one another and spread out on the surface; however, on the soft substrate, cells merge to form tissue aggregates [13, 24]. Guo *et. al* have observed this kind of behaviours in case of fibroblasts, epithelial cells and neonatal rat heart tissue. They found on soft substrate, cells form weak adhesions and are highly motile. Cells extend filopodia-like extensions that contract to bring the nearby cells into an aggregate [24]. Their findings show that cell-cell and cell-substrate mechanical interactions regulate tissue formation. It is observed that many cell types like fibroblasts, muscle cells exert contractile forces to the extracellular matrix (ECM) to probe its mechanical properties [25, 26]. Each active cell generates a deformation field in the medium that is sensed by the adjacent cells, leading to cell-cell mechanical interactions [14, 27]. Mechanical forces serve as cues for cellular rearrangements affecting tissue patterning and shapes in both animal and plant tissues [16, 28]. The mechanical signals are found to alter cell adhesion, migration, contractility, and many other activities and that in turn influence several physiological processes such as tissue morphogenesis, angiogenesis, myotube fusions, cancer metastasis, etc [16, 29–33]. Further, cells respond to not only the mechanical signals but also the chemical cues in the medium independent of the substrate mechanics. Many studies have shown cells can assemble and disassemble navigating the chemical gradients of chemoattractant or chemorepellants [34–36]. Chemotaxis provides an important route for long-range cell-cell communications and, as a result, facilitates cell aggregation and tissue growths [36–38]. Chemical signalling influences the motile cells to navigate according to the environmental changes and promotes aggregation by initiating cell-cell contacts. Further, it has been observed that motility mechanisms such as flagella, pili, gliding motion are controlled by the chemotaxis signal transduction [36]. The swarm formation in many species is found to be dependent on Chemotaxis. Myxococcus xanthus, a predatory bacterium in soil, coordinates its motility depending on spatiotemporal chemical gradients to swarm, predate and self-orgnize during fruiting bodies development [39]. Studies of social amoeba, in particular Dictyostelium discoidum, have shown the role of the chemotactic interaction in aggregation that leads to the formation of a multicellular organism [37, 40] There are several other examples of chemotaxis driven cellular assembly such as organization of cells into vascular networks driven by angiogenic stimulus, multicellular aggregate formation due to quorum sensing, wound healing, to name a few [36, 38, 41].

Thus, multiple pathways can promote the aggregation process; however, the difficulty lies in distinguishing the underlying mechanisms of cell-cell interactions that govern the cellular organization and lead to specialized tissue formation. Despite a multitude of attempts, our understanding is far from complete, and it remains a long-standing question in developmental biology. In this paper, based on a suitably constructed theoretical model, we probe how different pathways of cell-cell interactions affect the dynamics of aggregation from a seemingly random single cell population on a substrate. We investigate three distinct mechanisms of cellular interactions driven by physical cell adhesion contacts, mechanical interactions mediated through the extracellular matrix, and chemical signalling. Our analysis shows that the varying nature of cellular interactions strongly influences several properties of the aggregates, emerging tissue patterns, growth kinetics, and predicts a method to determine specific interaction pathways. We find, in the case of physical and mechanical interactions, the average domain size and the average mass of the growing clusters exhibit power law growth in time. In contrast, chemical signalling driven aggregation clearly deviates from this trend. Besides, the cluster size distributions can further characterize the distinct cellular growth processes. Our model also elucidates the effect of varying substrate stiffness on different aggregation pathways, as observed in experiments.

## THEORETICAL MODEL

We consider *N*_0_ cells are randomly deposited on a 2D lattice and interacting via one of the three pathways either by direct cell adhesion contacts, mechanical interaction, or chemical signalling. The detailed formulation is discussed in the following.

### (a) Aggregation due to physical cell adhesion contact

In many cell types, cell-cell adhesion, mediated by transmembrane junction proteins such as cadherins, play an important role to promote cellular assembly. Cells also form dynamic, filopodia-like extensions to interact with other cells when they come in close contact and bring them into an aggregate [24]. Experiments performed by Steinberg and collaborators demonstrated that cell aggregation and selective spreading of cells in a mixed cell population could occur simply due to the differences in the interfacial adhesion strength even in the absence of any differential chemical cues [42] which could be explained by Differential adhesion hypothesis (DAH). DAH provides an insight into such aggregation of cohesive cells and inspired many discrete cell models to understand various morphological structures during the development [20-–23, 43, 44]. We construct our model based on DAH, where motile and cohesive cells spontaneously adhere with each other to maximize the cohesive binding strength and minimize the cell-ECM interfacial energy of the system. DAH provides a connection between the tissue surface tension and the strength of adhesion between the cells constituting the tissue. In our simulation, the occupancy of each lattice site, *r* = (*i, j*), is specified by a type index, *σ_r_*, which we assume a value 1 if occupied by a cell and 0 for the medium. The total interaction energy of the system is given by, *E_p_* = ∑_<*r r′*>_ *J*(*σ_r_, σ_r′_*). Here, *J*(*σ_r_, σ_r′_*) is the interaction energy between two neighbouring pairs located at *r* and *r′*; the cell-cell interaction energy, *J*(1,1) = −*ϵ_cc_*; the cell-medium interaction energy, *J*(0,1) = *J*(1,0) = – *ϵ_cm_;* and the interaction between two medium sites is given by, *J*(0, 0) = – *ϵ_mm_* = 0. *ϵ_cc_* and *ϵ_cm_* denote the interaction strengths between cell-cell and cell-medium pairs. Now, *E_p_* can be rewritten separating the interfacial contribution and the bulk terms in the energy and given as

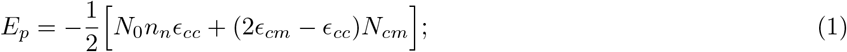

where *n_n_* is the significant nearest neighbours (in our model, *n_n_* = 8 on 2D substrate), and *N_cm_* denotes the total number of cell-medium bonds [45] (see supplementary). Cell aggregation or spreading occurs depending on the competition between the cell-cell cohesivity (*ϵ_cc_*) and the cell-substrate adhesivity (*ϵ_cm_*). Here, *γ_cm_* = (*ϵ_cc_*/2 – *ϵ_cm_*) denotes the cell-medium interfacial tension [4, 44–46]. If 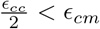, cells have the tendency to form aggregates and for 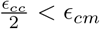, cells remain spread on the substrate.

### (b) Aggregation mediated by cell-substrate mechanical interactions

Several experiments show that cells build up a large number of focal adhesion contacts at the cell-matrix interface and exert contractile forces to the ECM to probe the mechanical properties of the surroundings [25]. The cellular contractile forces tend to deform the underlying substrate; cells can detect the matrix deformation created by the traction forces of the adjacent cells. The cell traction force, thus, provides matrix mediated cues for cell-cell communications and cells respond to it by actively regulating its position, orientations, migration, and various other activities [47–52]. Reinhart and coworkers have shown in their experiments that cells can detect and respond to the substrate strains generated by the contractile force of a neighbouring cell [14]. The cell response and also relative cell movements change depending on the varying substrate stiffness. Their findings show that cells communicate mechanically through the substrate, and the matrix mechanics foster tissue formation by promoting the formation of cell-cell contacts. The mechanosensing of individual cells influences the behaviour of the neighbouring cells, which translates into a collective organization of an ensemble of cells in a compliant environment and promotes cellular assembly and tissue development [14, 25, 27]. In a coarse-grained approach, cells can be modelled as contractile force dipoles or contractile disks that generate elastic deformation of the medium [25, 53]. Following earlier studies [53], we consider the cells as contractile disks on a 2D semi-infinite elastic substrate. In our model, we consider isotropically contractile cells, hence restrict our analysis to the case of radial forces. The interaction energy between two active contractile disks is given by, 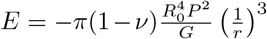, where *G* is the elastic modulus, and *v* is the Poisson’s ratio of the substrate. *P* denotes the force magnitude, *R*_0_ is the radius of the cell disk, and *r* is the distance between the cells [53]. In our case of a linear elastic medium, since the strain fields superimpose, the elastic interaction energy of a system of *N*_0_ cell can be expressed as,

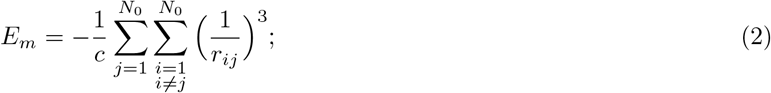

where, *r_ij_* is the distance between *i^th^* and *j^th^* cell and for a given cell type, *c* is proportional to the elastic modulus of the medium (other factors including the factor (1/2) required to avoid the double counting are absorbed within the constant *c*). The interaction energy depends on the distance, *r_ij_*, between the cells. Since the interaction energy is negative, isotropically contractile, active cells would be attracted to each other, and the interaction strength decays with the increase in the distance between the cells.

### (c) Aggregation driven by chemical signalling

Several experiments suggest that motile chemotactic cells sense and respond to the changes in their chemical environments and drive themselves in favour of the growth process [34, 36]. Besides, many cell types, bacteria, slime molds, among others, secrete chemicals to interact with each other via chemosignaling and drive the aggregation process [35, 37]. Chemotaxis provides pathways for long-range cellular communication and develops transient cell-cell contacts. Eventually, cells following the chemical cues navigate themselves to the most favourable niches and form multicellular aggregates [36]. Keller and Segel [37] have developed a mathematical formulation of cellular aggregation driven by chemotaxis. They have proposed that cells move towards a relatively higher concentration of chemicals secreted by the cells viewing it as drift-diffusive motion in the direction of chemotactic attraction. Based on this framework of chemotaxis model, Lee *et al.* [54] have further studied the mechanisms of interactions between chemotactic cells. They have considered that individual cells produce chemoattractant that diffuses in space and decays exponentially with time. Due to diffusion and degradation, the released chemoattractants form a spatial gradient around the cell leading to the cell-cell attraction. Based on these earlier studies, we present a minimal model to incorporate the long-range attraction of cells due to chemical signalling. In our model, we consider an effective cell-cell interaction energy due to chemical signalling of the form [54, 55],

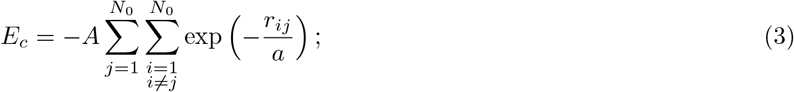

where *r_ij_* is the distance between a pair of cells, *i* and *j*, and *A* is the strength of the chemotactic effect. Here, *a* signifies the spatial range of the chemoattractant determined by 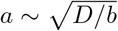, where *D* is the diffusion coefficient and *b* is the decay rate of the chemical signals. Moreover, chemotactic speed is found to be sensitive to the substrate stiffness. Saxena *et. al.* have performed experiments to demonstrate the collective influence of physical and chemical cues in chemotactic cell migration [56]. They varied the substrate stiffness and observed that the chemotactic speed is scaled inversely with substrate stiffness. It is found that weaker adhesion of cells and higher protrusion rate on softer substrates help faster chemotaxis. In our model, the larger range of the chemoattractant, *a*, helps the aggregation process [as shown in Fig. S1(c) in supplementary]. As on softer substrates, chemotactic speed is faster and promotes the aggregation process; in our model, it is tuned by the parameter *a*.

### Numerical algorithm

At each simulation step, all *N*_0_ cells are picked up one by one at random and given a probability to move to any of its empty eight neighbouring sites following the interaction energy for three different pathways as given by Eqs. (1), (2), and (3) respectively. Each simulation step is considered as one unit time step evolution. We calculate the energy, *E*_0_, of the chosen cell in its current location and also its energy, *E_n_*, in the possible new location. If *E_n_* ≤ *E*_0_, then the move is readily accepted as it lowers the energy of the system. However, if *E_n_* > *E*_0_, then a random number, *R*, is generated from an uniform distribution over the interval (0,1) and the cell moves to the higher energy site with a probability such that *R* < exp(–Δ*E*), where Δ*E* = *E_n_* – *E*_0_.

## RESULTS

We investigate the initiation, growth, and temporal evolution of the cellular aggregates governed by three mechanisms of cell-cell interactions due to direct cell adhesion contacts, matrix mediated mechanical interactions, and chemical signalling. Following these three distinct pathways, cells reorganize their positions by breaking existing bonds and making new ones. They also move and merge with other cells leading to growing aggregates. Our simulations show that the growth kinetics strongly depend on the nature of the cell-cell interactions. It is also sensitive to the rigidity of the substrate, as seen in experiments. We have investigated the dynamics for a wide range of parameter values, here, we present some representative simulation results considering *N*_0_ = 1000 cells placed on a lattice of size, *L* = 200.

Figures 1(a)-(b) present snapshots of the temporal evolution of cell aggregation driven by physical adhesion contacts. In this case, competition between cell-cell cohesivity, *ϵ_cc_*, and cell-substrate adhesivity, *ϵ_cm_*, determines the aggregation process as described by Eq. 1. On a stiffer substrate, cells build up mature adhesion contacts, so cell-substrate adhesion is stronger, and cells remain spread out. On the other hand, a softer substrate where cell cohesivity dominates is more favourable for cell aggregation. Figure 1 shows one such cellular aggregation where the interaction strengths are chosen as *ϵ_cc_* = 10.0 and *ϵ_cm_* =0.1. Due to strong cell-cell binding affinity (*ϵ_cc_*), when the cell finds another neighbouring cell, sticks to one another, and form nascent clusters as shown in Fig. 1(a). These clusters gradually grow with time due to the merging of other cells in the vicinity and give rise to larger aggregates, as seen from Fig. 1(b). It is worth noting that if the clusters are allowed to diffuse as a whole, they would merge and form bigger aggregates.

**FIG. 1.**
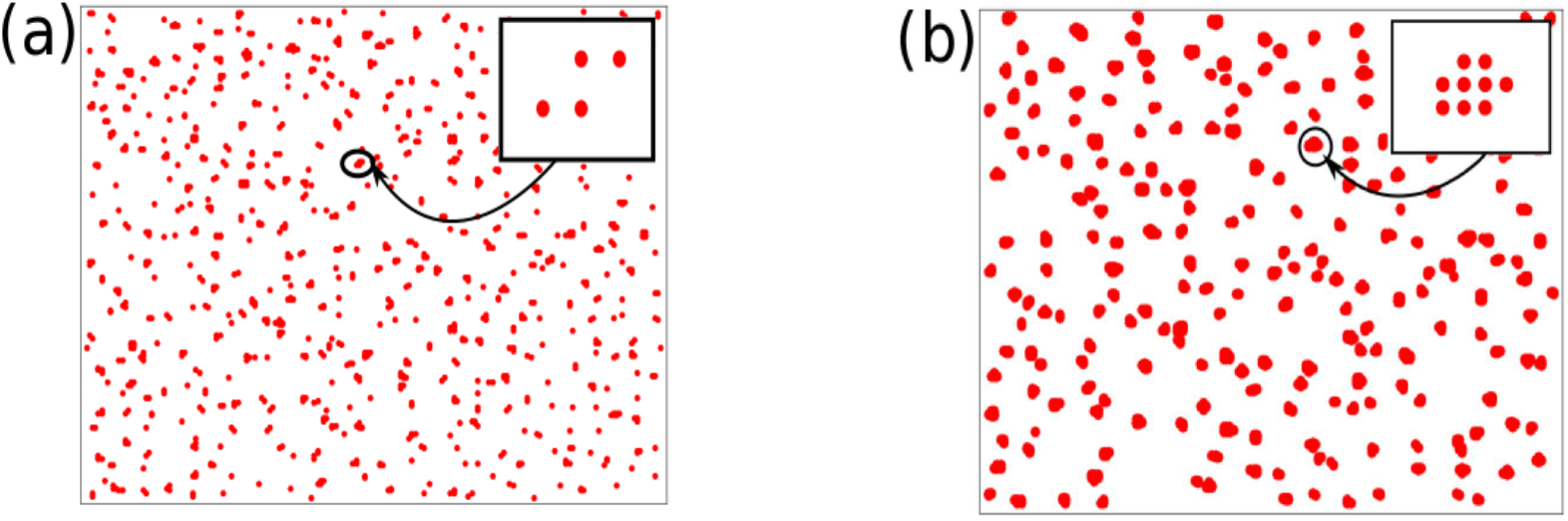
Time evolution of cellular aggregation due to physical cell adhesion contacts. Snapshots of growing aggregates at time, (a) *t* = 100 and (b) *t* = 1000. The inset shows one aggregate structure. Here, the value of *ϵ_cc_* = 10 and *ϵ_cm_* =0.1, representing a softer substrate that promotes cell aggregation.

On the other hand, in the case of mechanical interactions, cells interact with each other mediated through the elastic substrate, which gives rise to an effective attraction among cells. As time progresses, cells start to migrate to reach the lowest energy configuration given by Eq. 2 and the clusters grow with accumulations of more cells. Figures 2(a)-(b) show the snapshots of such growing clusters. Moreover, as the interaction energy varies as *ϵ_m_*(*r*) ~ 1/*r*^3^, the cellular attraction decays with the increase in the distance, *r*, between cells; therefore, the domain size of the clusters grow up to a certain value. Results presented here are for a smaller value of *c* = 0.1 that represents a softer substrate, as c is proportional to the elastic modulus of the substrate. However, with higher values of *c*, *i.e.,* on stiffer substrates, cells remain spread out (shown in supplementary).

**FIG. 2.**
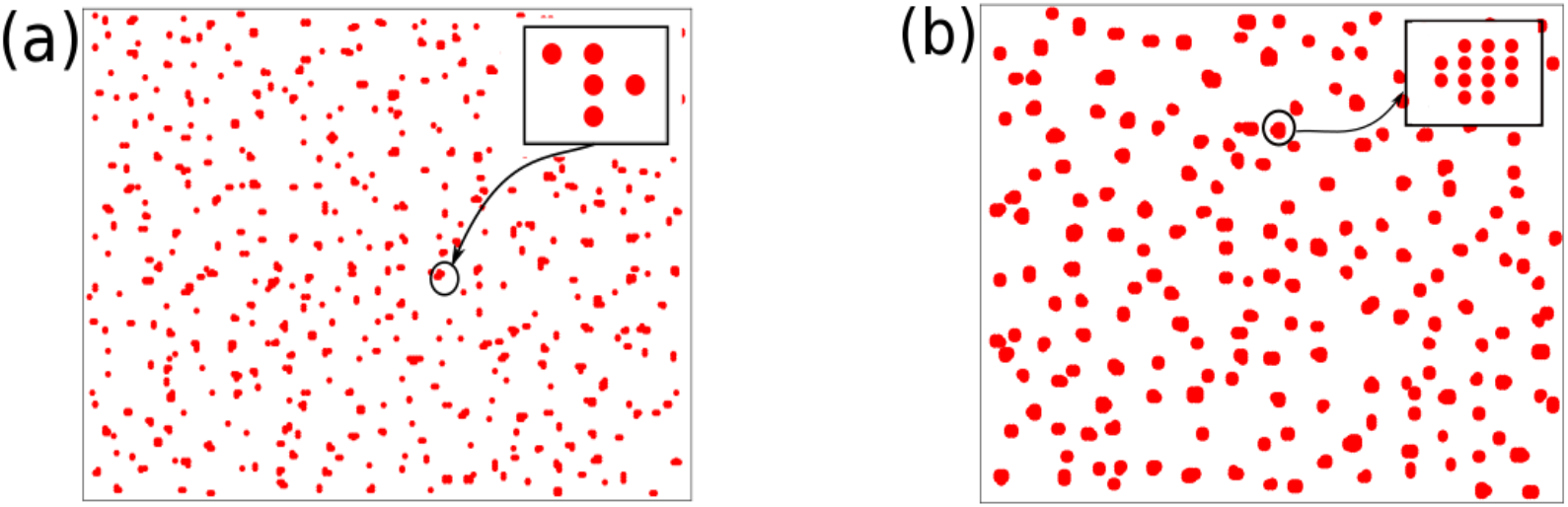
Time evolution of aggregates formation mediated by cell-substrate mechanical interactions. Here, the snapshots are at time, (a) *t* = 100 and (b) *t* = 1000. One aggregated structure is shown in the inset. The value of *c* = 0.1 representing a softer substrate and favourable for cell aggregation.

Next, Fig. 3 presents the temporal evolution of formation of aggregates due to chemical signalling. Our study shows that the spatial range, *a,* of the chemotactic signals greatly influences the formation and structure of the cellular aggregates (as described by Eq. 3). For a smaller range, *a* = 10, of the diffusive chemoattractants, as the interaction is limited to the adjacent cells, many small clusters are formed, as shown in Figs. 3(a)-(c). However, at large values of *a* = 100, due to long-range communications, all cells come together to form a bigger aggregate, as seen from Figs. 3(d)-(f) (keeping the strength of chemotactic signal, *A* = 10). Moreover, as found in experiments that chemotaxis happens faster on the softer substrate [56], in our simulations, the parameter *a* could capture such dependence of the aggregation process on varying substrate stiffness. Since *a* varies with the diffusion of the chemicals as, 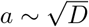; thus, faster the diffusive speed, higher the value of *a*. This, in turn, implies large aggregation is favourable on a softer substrate which is in agreement with experiments.

**FIG. 3.**
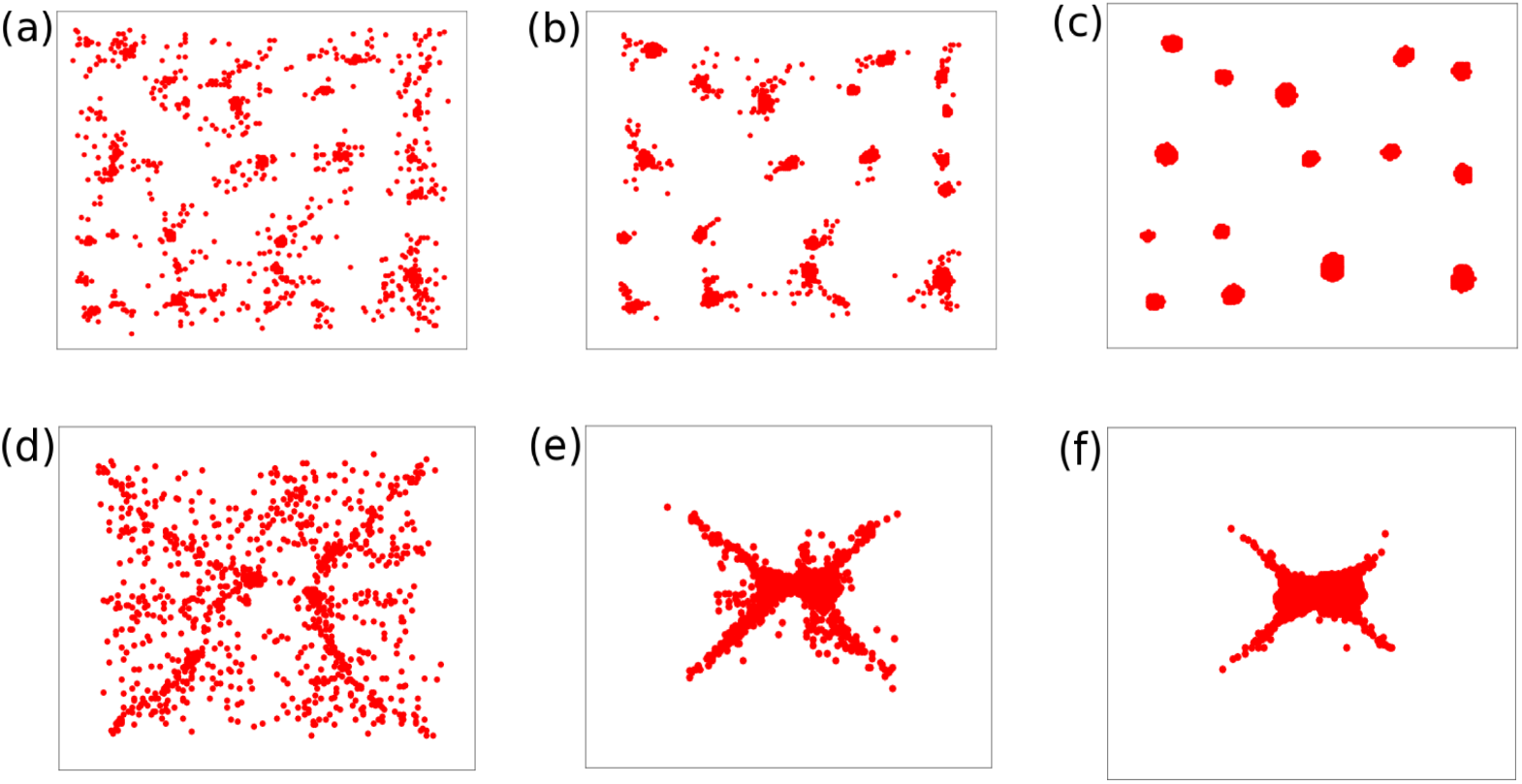
Time evolution of aggregation of cells driven by chemical signalling. (a)-(c) present snapshots of growing aggregate at different time, *t* = 100, 300, and 1000 respectively for *a* = 10, representing a smaller spatial range of chemoattractant; whereas (d)-(f) show cluster formation for a longer range *a* = 100 at the same time snapshots.

### Rate of cell aggregation

We now investigate the rate of number of clusters formation mediated by different interaction pathways. In our simulation, we start with *N*_0_ number of single-cell clusters. As time progresses, the cells migrate, join together and form small clusters depending on the nature of the cell-cell interactions. Thus, as the aggregates grow, the total number of clusters decreases with time. Figure 4 shows the time evolution of number of clusters, *N*(*t*), normalized by *N*_0_. As shown in the figure, in the case of physical and mechanical interactions, the number of clusters decreases faster as cells merge together rapidly due to local cell-cell attractions and form smaller domains of aggregates. On the contrary, in chemical signalling, the rate of cluster formation is sensitive to the spatial range of interactions (*a*) among cells via diffusive chemicals, and cells form much bigger aggregates. We have further investigated the effect of cell density, *ρ* = *N*_0_/(*L* × *L*), on the aggregation process. We find that with the increase in density, the growth process gets faster as migrating cells readily find each other in their immediate vicinity and the cluster size also grows bigger due to the availability of more cells.

**FIG. 4.**
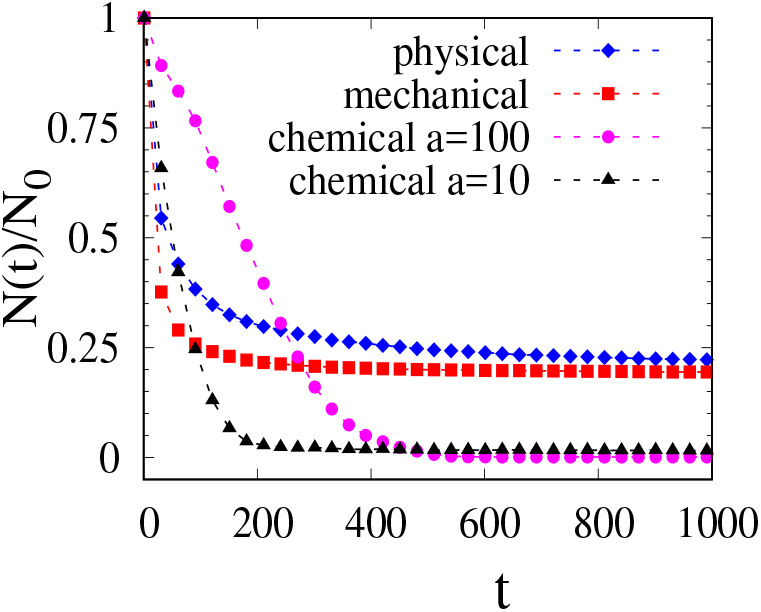
Time evolution of number of clusters formation due to three distinct cell-cell interaction pathways mediated by physical 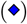, mechanical 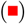, and chemical signalling with a smaller range, *a* = 10 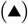, and a longer range, *a* = 100 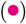.

### Power law growth of the aggregates

To distinguish the underlying mechanisms of different interaction pathways further, we analyse the growth kinetics of the aggregation process. In physical and mechanical interactions, the average domain size of the clusters, *l*, exhibits a power law growth with time as, *l*(*t*) ~ *t^α^*. Figure 5(a) shows *l* as a function of time, *t*, on a log-log scale. We also investigate the average mass, *m*(*t*), of the growing clusters. In our model, each cell is considered unit mass; hence, the aggregate’s mass is calculated by the total number of cells in that aggregate. Time evolution of the average mass of the clusters also demonstrate a power law behaviour, *m*(*t*) ~ *t^β^*, as shown in Fig. 5(b). As time progresses, the average cluster size increases due to the joining of more and more cells, and thus, the clusters grow bigger. After some time, gradually, the growth saturates and reaches a steady-state configuration depending on the range of cellular interactions. However, chemotaxis driven aggregation mechanism keeping *A* =10 and *a* = 100 distinctly deviates from this power law growth, as shown in the insets of Figs. 5(a) and (b). The power law exponents, *α* and *β*, turn out to be higher in the mechanical interaction (*c* = 0.1) of cells. Also, the growth process is faster, and the aggregates are formed bigger compared to physical adhesion (*ϵ_cc_* = 10, *ϵ_cm_* = 0.1). Interestingly, we find that substrate stiffness has a stronger influence on cellular aggregation mediated by cell-substrate mechanical interactions compared to the direct contact adhesion route. Besides, the power law exponent does not vary significantly with change in the cell density, system size, or the substrate stiffness in the softer regime (that favours cell aggregation as shown in supplementary); thus, signifies that the specific growth behaviour is intrinsic to the underlying pathways of cell-cell interactions. It is noteworthy that power law growth of cluster domains has been observed in many active and passive systems that exhibit clustering phenomena such as in the self-propelling motion of active particles, phase separation in binary mixtures, droplet growth, among others [57–60]. The power law exponents depend on the mechanisms of transport, diffusion and coalescence of the clusters, temperature and the type of the patterns. Lifshitz-Slyozov [61] have shown such power-law growth kinetics in systems where growth occurs via evaporation-condensation mechanisms and the exponent turns out to be *α* = 1/3. Further, Binder and Stauffer [59] have also shown power-law domain growth in the case droplets or clusters diffuse and coalesce following collisions. Considering this mechanism, the evolution of the clusters can be described by the kinetic equation, 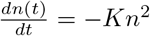 [58, 62, 63]. Here *n* is the cluster density, *l* is the size of the cluster (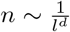, *d* is the spatial dimension), and *K* is the rate constant. Now, treating *K* as a constant, the equation predicts the growth law as *l*(*t*) ~ *t^α^* with the exponent *α* = 1/*d*. However, *K* depends on both size, *l*, and the diffusivity, *D* of the clusters and thus, it modifies the growth rate. The growth exponent, *α*, changes depending on the system, the interaction mechanisms, and the mobility or diffusivity of the clusters. In our study, clusters are not allowed to diffuse; here, the power law growth arises due to the cell movements governed by specific cell-cell interactions in the aggregation process. In the case of physical adhesion, cell motility arises due to the local interaction of neighbouring cells via adhesion contacts, and in the case of mechanical interaction, cellular interaction strength decreases with the increase in the distance, *r*, between the cells as 1*/r*^3^. These short-range interactions result in lower cell mobility and slower cluster growth; thus, the growth exponent turns out to be smaller. We also study the power law growth with the Kinetic Monte Carlo method to see how the dynamics evolve in the physical description of time. We find that the qualitative behaviours of the growth kinetics of three different pathways remain the same with both the simulation methods (presented in supplementary).

**FIG. 5.**
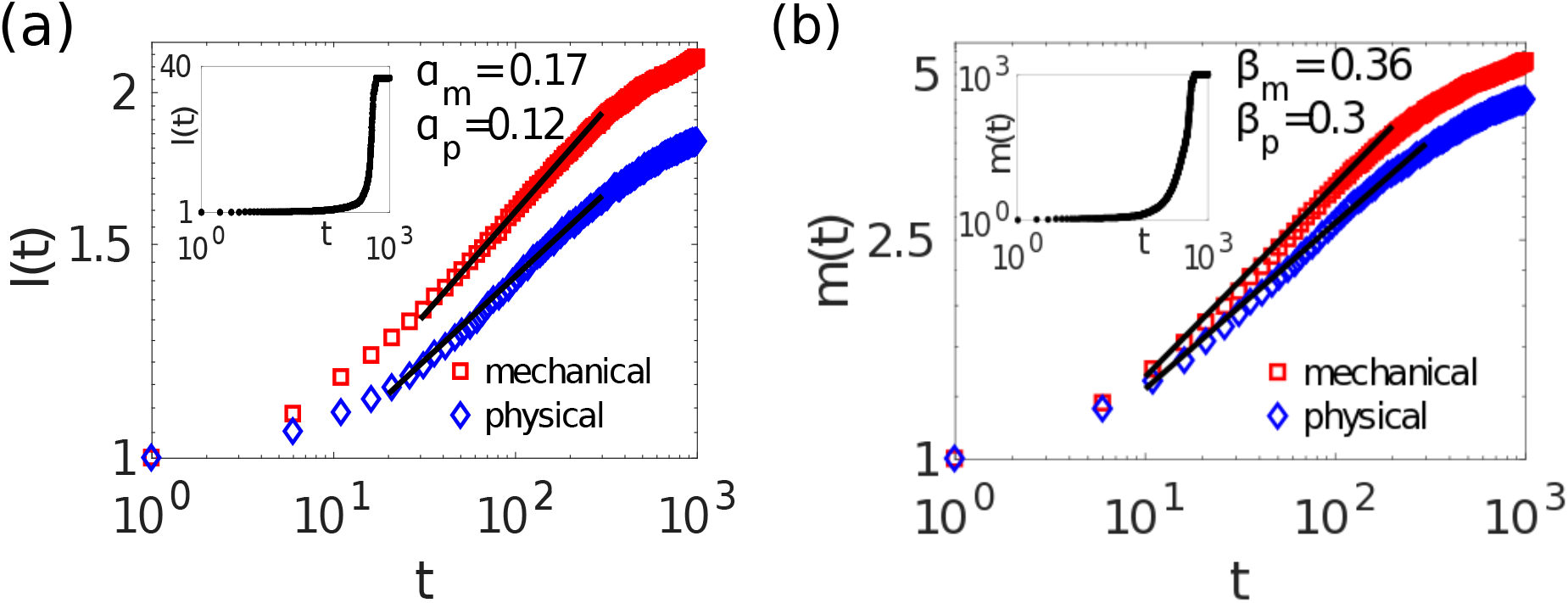
Time evolution of the characteristic growth of the aggregates under physical 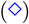, mechanical 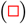, and chemical 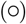 interactions. (a) The average cluster size, *l,* as a function of time, *t*, on a log-log plot. The solid lines show a power law growth with the exponent value, *α_p_* = 0.12 and *α_m_* = 0.17 for physical and mechanical interactions respectively. (b) Log-log plots of the average mass, *m*, of the clusters as a function of time with the power law exponents, *β_p_* = 0.3 and *β_m_* = 0.36 for physical and mechanical interactions respectively. The insets show the growth due to chemical interaction.

### Cluster size distribution

We further characterize the aggregation kinetics by analysing how the distribution of cluster size changes as time progresses. The cluster size distribution function, *p*(*n, t*), denotes the fraction of clusters of size *n* at time, *t*. Here *n* is the number of cells in a cluster of size *n*. Since there is fluctuations in cluster size variation, we consider a comparatively smooth distribution function, namely, complementary cumulative distribution (CCD) function given by *P*(*n,t*) = 1 – ∑_*i*<*n*_ *p*(*i,t*), where *P*(*n,t*) denotes the probability of occurrence of number of clusters with size equal or greater than *n* at time *t*. Figures 6(a)-(c) show the evolution of the CCD function, *P*(*n,t*), at different times for three leading aggregation mechanisms. For physical [Fig. 6(a)] and mechanical [Fig. 6(b)] interactions, the distributions can be fitted by stretched exponentials as *P*(*n,t*) = *A*exp[ – (*n/n_s_*)^*τ*^]. The exponent *τ* determines the fatness of the tail of the stretched exponential distribution; with *τ* ≤ 1, the smaller the *τ* value, the fatter is the tail, and *n_s_* is a characteristic scale from which all moments can be evaluated [64, 65]. The borderline *τ* = 1 corresponds to the usual exponential distribution. For *τ* smaller than 1, the distribution presents a clear curvature in a log-log plot indicating a clear deviation from the power law scaling. Moreover, the smaller is *τ*, the greater is the linear behaviour in a log-log plot.

**FIG. 6.**
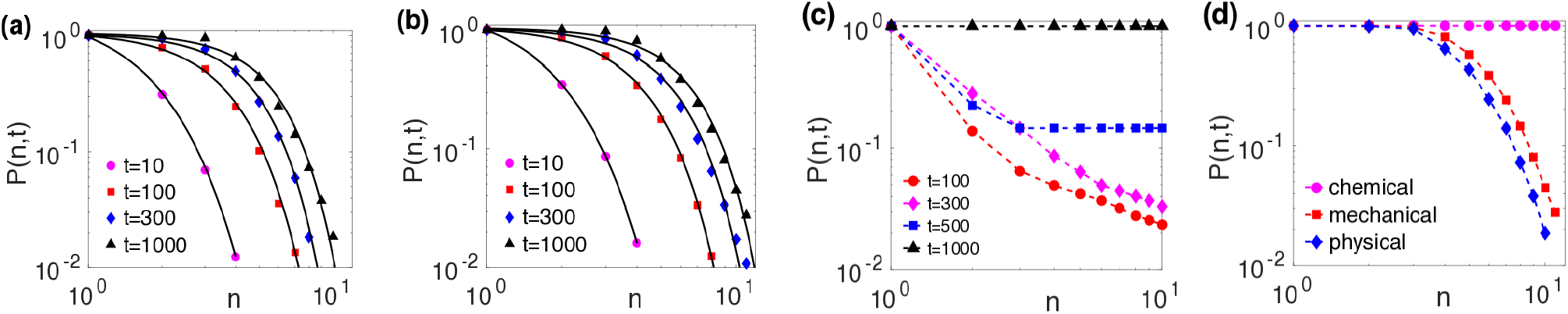
Log-log plots of cumulative cluster size distribution (CCD) function, *P*(*n,t*), under three different cellular communication pathways. (a)-(b) Show the evolution of CCDs for physical and mechanical interactions at different time, *t*. The solid black lines correspond to stretched exponential fittings to the simulation data. (c) Time evolution of CCDs for chemotaxis driven aggregation. It does not exhibit stretched exponential decays. (d) Comparison of distribution functions for three interaction pathways at steady state.

The distribution function, initially with the formation of small clusters, shows a sharper tail with a higher value of *τ*; with time, as the clusters grow in size, it exhibits a neat curvature giving rise to more fat-tailed stretched exponential decays. In the steady state, the mechanical interaction exhibits a more fatter tail compared to the physical interaction indicating the formation of larger clusters (the values of *τ* at *t* = 10, 100, 300, 1000 for physical interaction are 0.66, 0.5, 0.46, 0.43 and that for mechanical interaction are 0.7, 0.5, 0.42, 0.4 respectively). The stretched exponential cluster size distributions have also been observed in experimental studies on aggregation kinetics of some normal cell lines of epithelial and mesenchymal origin and malignant cell lines [64]. On the other hand, for chemotactic interactions, the distribution function does not exhibit stretched exponential decay, as seen from Fig. 6(c). The log-log plots of the distribution function exhibit a linear region at the early stages; however, as time progresses, due to long-range interactions, one big aggregate or a few large aggregates are formed.

## DISCUSSION

In conclusion, our work provides many insights in distinguishing the underlying mechanisms of cell-cell interaction pathways in the aggregation process, which were hitherto unexplored. Our study delves deeper into the growth kinetics of three leading cellular interaction mechanisms and shows that the emergence of power law growth and the stretched exponential cluster size distributions uniquely mark the differences between different cell-cell communication pathways. Both physical cell adhesion and mechanical interactions exhibit power law growth in average domain size and the mass of the clusters and also display stretched exponential cluster distributions. However, the growth process turns out to be faster under matrix mediated cell-cell mechanical interactions compared to the physical contact adhesion route. On the other hand, chemotaxis driven aggregation deviates distinctly from these trends. Moreover, the value of the power law exponents turns out to be pretty robust, with the variation in cell density, system size, and substrate rigidity showing an intrinsic characteristic of the cell-cell interaction mechanisms. Further, our study shows that substrate stiffness has a stronger influence on cellular aggregation mediated by cell-substrate mechanical interactions compared to the direct contact adhesion route. As shown in the supplementary, the power law exponents do not change with the variation in the substrate stiffness in the case of physical adhesions, and the growth curves almost collapse into a single curve. On the other hand, in the case of mechanical interactions, the growth process turns out to be faster with the decrease in substrate stiffness. The stretched exponential cluster size distributions, as predicted by our model, have also been observed in experiments for some normal and malignant cell lines [64]. As stretched exponential distributions have been found in many natural systems, it indicates a universal cluster size distribution in aggregation kinetics [65]. Our findings offer powerful tools that can further be tested in experiments for various cell types and varying substrate stiffness to identify the underlying pathways of cell-cell communications. As tissue development is a complex process, our study based on a simple, generic model of active cellular aggregation is envisaged to pave the way for further extended studies to understand the governing pathways of cellular organizations that would not only be immensely beneficial in diverse areas of developmental biology but also help fine-tune therapeutic approaches of several diseases, cancer metastasis, and in the field of artificial tissue engineering.

## AUTHOR CONTRIBUTIONS

R.D. conceptualized the research. D.M. carried out all simulations. D.M. and R.D. analysed the data and wrote the manuscript.

## ACKNOWLEDGEMENTS

We are grateful for discussions with Prof. Samuel Safran and the financial support from SERB, Grant No. SR/FTP/PS-105/2013, DST, India.

